# Discovery of a *Streptococcus pneumoniae* serotype 33F capsular polysaccharide locus that lacks *wcjE* and contains a *wcyO* pseudogene

**DOI:** 10.1101/308767

**Authors:** Sam Manna, Eileen M. Dunne, Belinda D. Ortika, Casey L. Pell, Mike Kama, Fiona M. Russell, Tuya Mungun, E. Kim Mulholland, Jason Hinds, Catherine Satzke

## Abstract

**Objectives:** As part of large on-going vaccine impact studies in Fiji and Mongolia, we identified 25/2750 (0.9%) of nasopharyngeal swabs by microarray that were positive for *Streptococcus pneumoniae* contained pneumococci with a divergent 33F capsular polysaccharide locus (designated ‘33F-1’). We investigated the 33F-1 capsular polysaccharide locus to better understand the genetic variation and its potential impact on serotyping results.

**Methods:** Whole genome sequencing was conducted on ten 33F-1 pneumococcal isolates. Initially, sequence reads were used for molecular serotyping by PneumoCaT. Phenotypic typing of 33F-1 isolates was then performed using the Quellung reaction and latex agglutination. Genome assemblies were used in phylogenetic analyses of each gene in the capsular locus to investigate genetic divergence.

**Results:** All ten pneumococcal isolates with the 33F-1 *cps* locus typed as 33F by Quellung and latex agglutination. Unlike the reference 33F capsule locus sequence, DNA microarray and PneumoCaT analyses found that 33F-1 pneumococci lack the *wcjE* gene, and instead contain *wcyO* with a frameshift mutation. Phylogenetic analyses found the *wzg, wzh, wzd, wze, wchA, wciG* and *glf* genes in the 33F-1 *cps* locus had higher DNA sequence similarity to homologues from other serotypes than to the 33F reference sequence.

**Conclusions:** We have discovered a novel genetic variant of serotype 33F, which lacks *wcjE* and contains a *wcyO* pseudogene. This finding adds to the understanding of molecular epidemiology of pneumococcal serotype diversity, which is poorly understood in low and middle-income countries.

## Introduction

*Streptococcus pneumoniae* (the pneumococcus) is a Gram-positive pathogenic bacterium and a leading cause of community-acquired pneumonia [1]. Pneumococci are classified by serotype, defined by an antigenically-distinct polysaccharide capsule. Capsule biosynthesis is encoded by the capsular polysaccharide (*cps*) locus within the pneumococcal genome. High levels of genetic diversity within this locus has resulted in over 90 pneumococcal serotypes described to date.

The pneumococcal capsule is the target for currently licensed vaccines, which only include a subset of serotypes. Although pneumococcal conjugate vaccines (PCVs) have been successful in reducing carriage and disease caused by the targeted serotypes, a rise in carriage and disease caused by serotypes not included in these vaccines is commonly observed (serotype replacement) [2,3]. To precisely monitor vaccine impact and disease surveillance, accurate tools for pneumococcal serotyping are required.

Molecular approaches to serotyping pneumococci rely on existing knowledge of *cps* loci. Data on pneumococcal *cps* loci from low- and middle-income countries (LMICs) are relatively limited, which can impact serotyping results. For example, we recently described a novel genetic variant of pneumococcal serotype 11A in Fiji. Genetically, the *cps* locus of these isolates is most closely related to the 11F *cps* locus, with only a few minor nucleotide changes resulting in the production of 11A capsule [4].

Among the replacing serotypes post-PCV introduction, serotype 33F has become a concern world-wide. Serotype 33F is commonly reported among the predominant serotypes not included in PCVs causing invasive disease following vaccine introduction [5–7]. The increased invasive disease caused by serotype 33F has warranted its inclusion in two new vaccine formulations, which are in development by Merck [8]. In this study, we describe a novel 33F *cps* locus identified in Fiji and Mongolia by investigating the genetic basis of the variation in this locus and the potential impact this may have on serotyping results.

## Materials and Methods

### Nasopharyngeal swab collection and screening for pneumococci

As part of ongoing programs in the Asia-Pacific region measuring pneumococcal vaccine impact, nasopharyngeal swabs from healthy participants in Fiji, and children diagnosed with pneumonia in Mongolia were collected in accordance with WHO recommendations [9]. Ethical approval for the study in Fiji was granted from the Fiji National Research ethics review committee and The University of Melbourne Human research ethics committee. Ethical approval for the study in Mongolia was granted from the ethics committee associated with The Ministry of Health in Mongolia and the Royal Children’s Hospital in Melbourne. Written consent for study participants was provided by parents/guardians. Following collection, the swabs were placed in 1 ml skim milk, tryptone, glucose, and glycerol media [10] and stored at −80°C. Samples were screened for the presence of pneumococci by conducting quantitative PCR (qPCR) on DNA extracted from 100 μl aliquots of the swabs using the pneumococcal *lytA* gene as a target as previously described [11].

### Molecular serotyping by microarray

Molecular serotyping of pneumococci was performed by DNA microarray. An aliquot of the nasopharyngeal swab was inoculated onto Horse Blood Agar supplemented with gentamicin (5 μg/ml), to select for pneumococci, and incubated overnight at 37°C with 5% CO_2_. For plates with α-hemolytic growth, the bacterial growth was collected using 1 ml PBS, pelleted by centrifugation and stored at −30°C. DNA was extracted from thawed bacterial pellets using the QIAcube HT with the QIAamp 96 DNA QIAcube HT Kit (Qiagen) with the inclusion of a pre-treatment lysis step whereby 180 μl lysis buffer (20 mM TrisHCl, 2 mM EDTA, 1% Triton X-100, 2 mg/ml RNase A, 20 mg/ml lysozyme) was added to the bacterial pellet and incubated at 37°C for 60 min. The remaining extraction procedure was as per the manufacturer’s instructions. This DNA was then used for microarray as described previously [12]. In brief, 200 ng of DNA was labelled with Cy3 or Cy5 using the Genomic DNA ULS Labeling Kit (Agilent Technologies) and incubated at 85°C for 30 min. The labelled pneumococcal DNA was incubated with Senti-SPv1.5 microarray slides (BUGS Bioscience) overnight at 65°C rotating at 20 rpm. Microarray slides were washed, scanned, and analyzed using the Agilent microarray scanner and feature extraction software. Serotype calls were analyzed by Senti-NET software (BUGS Bioscience) using Bayesian-based algorithms.

### Bacterial isolates

The *S. pneumoniae* isolates used in this study (Table 1) were purified from ten nasopharyngeal swabs containing 33F-1 from Fiji and Mongolia on selective media as described above. Isolates were confirmed as *S. pneumoniae* with microarray and whole genome sequencing.

**Table 1.**
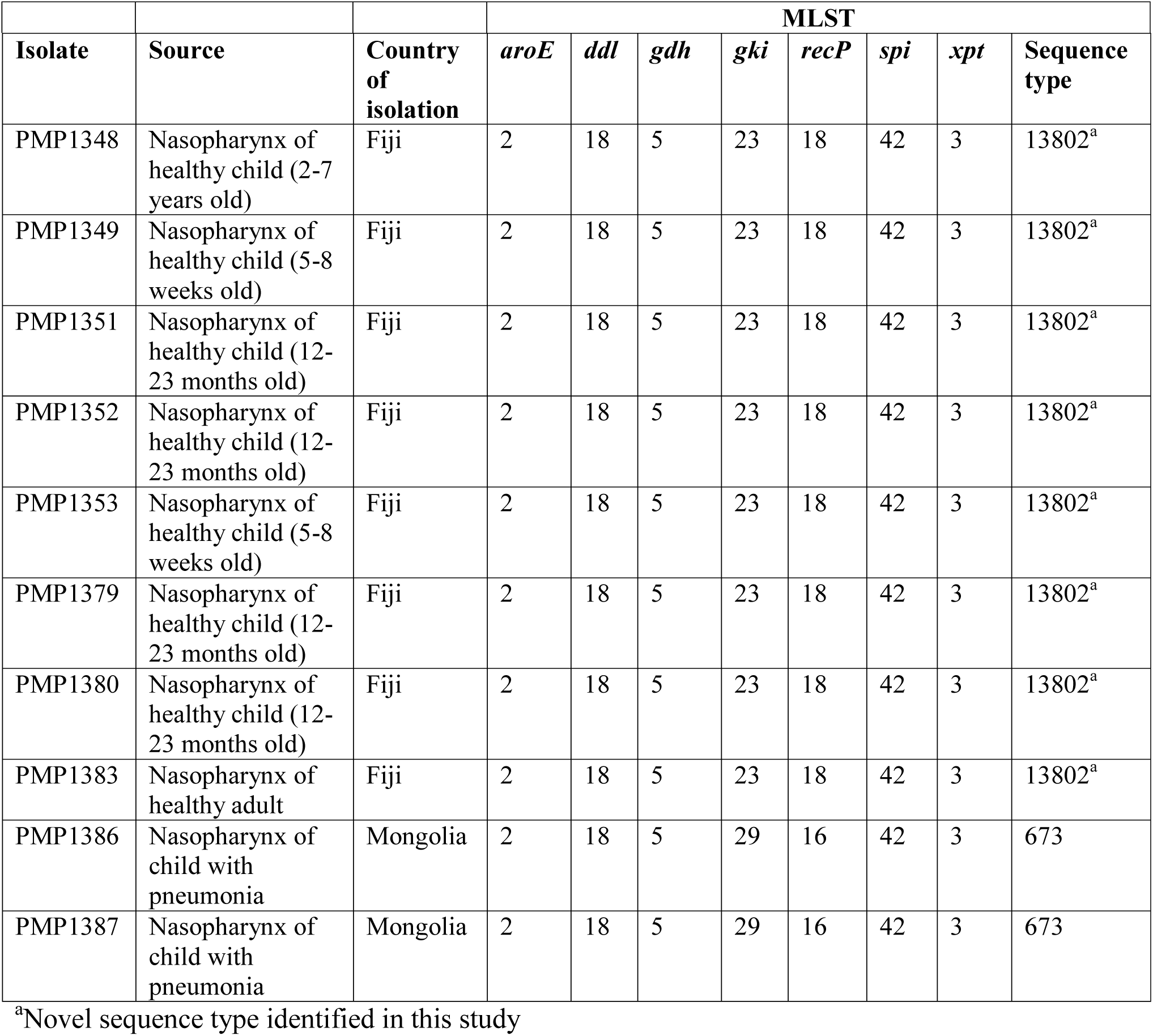
Pneumococcal 33F-1 isolates used in this study.

### Whole genome sequencing and molecular typing

For whole genome sequencing, DNA was extracted from pure cultures using the Wizard SV genomic DNA purification system (Promega) with some modifications. Briefly, pneumococcal cultures were pre-treated with a lysis solution containing 5 mM EDTA, 3 mg/ml lysozyme and 37.5 μg/ml mutanolysin in TE buffer and incubated at 37°C for 2 h. Proteinase K was added to a final concentration of 1 mg/ml and samples were incubated at 55°C for 1 h. Following incubation, 200 μl of nuclear lysis buffer and 5 μl of RNase (final concentration of 40 μg/ml) were added and samples were incubated at 80°C for 10 min. The remaining extraction procedure was performed as per the manufacturer’s instructions. Eluted DNA was sequenced in 2 × 300 bp paired end reads on the MiSeq platform. Using the Geneious 11.0.4 software package [13], sequence reads were trimmed with BBDuk and *de novo* assembled using SPAdes. The capsule loci were annotated within Geneious using a database consisting of capsule loci from the 90 serotypes described by Bentley et al. [14]. Sequence reads were also used for molecular typing with PneumoCaT [15].

### Sequence analysis

Pairwise alignments were using either MUSCLE or Clustal Omega. Phylogenetic analyses were performed for each 33F-1 *cps* gene using MEGA 7 [16]. For each gene, the phylogenetic analysis included a representative 33F-1 sequence as well as homologues from all other serotypes containing that gene as described by Bentley et al. [14], where Genbank accession numbers are provided. DNA sequences were aligned using MUSCLE and the alignments were used to generate maximum likelihood trees based on the Tamura-Nei model. Phylogenetic relationships were statistically analyzed by bootstrapping (1000 replicates). The 33F-1 *cps* loci have been deposited in Genbank (accession no. MH256127, MH256128, MH256129, MH256130, MH256131, MH256132, MH256133, MH256134, MH256135, MH256136).

### Quellung and latex agglutination serotyping

Quellung serotyping was performed as described previously [17]. A saline suspension of pneumococci was prepared from an overnight culture. Using an inoculation loop, 1 μl was placed on a microscope slide and mixed with 1 μl of antisera from the Statens Serum Institut (SSI) (http://www.ssi.dk/ssidiagnostica). The sample was then viewed under the microscope (x400 magnification). A positive reaction was defined as an enlargement or ‘swelling’ of cells, with serotype call based on the reaction profile with each typing sera. For latex agglutination, latex reagents were prepared with SSI typing sera [18] and testing performed as previously described [19]. The bacterial suspension and latex reagent (10 μl of each) were mixed on a glass slide. The slide was then incubated on an orbital shaker for 2 min at ~140 rpm. A positive reaction was defined by the presence of visible agglutination.

## Results

In our studies evaluating pneumococcal vaccine impact in Fiji and Mongolia, we have used DNA microarray as a molecular approach to serotype pneumococci contained within nasopharyngeal swabs. DNA microarray uses 15,000 oligonucleotides that are spotted onto glass slides and recognize each capsule gene from the 90+ serotypes. Labelled pneumococcal DNA is allowed to hybridize to the oligonucleotides so that pneumococcal serotype can be inferred. From 2750 swabs that contained pneumococci 25 (0.9%) contained pneumococci that typed as ‘33F-like’ (hereby referred to as ‘33F-1’). Ten of these samples were selected and the 33F-1 pneumococci were isolated for further analysis (Table 1).

Compared to the expected results for serotype 33F, microarray reported the *wciG, glf* and *wcjE* genes in the nasopharyngeal swabs containing these isolates as ‘absent/divergent’. In addition, the *wcyO* gene was also detected, which has not been reported in the serotype 33F *cps* locus previously. To investigate the impact of the divergent 33F-1 *cps* locus on other molecular approaches to serotyping, we sequenced the genomes of all ten isolates and ran the sequence reads through the PneumoCaT pipeline [15]. PneumoCaT uses *wcjE* to differentiate 33A from 33F, as this gene contains a frameshift mutation in 33F, resulting in a lack of WcjE-mediated O-acetylation of the 33F capsular polysaccharide [20]. Consistent with microarray, PneumoCaT typed all isolates as 33F and was unable to detect *wcjE*. Phenotypic serotyping methods (Quellung and latex agglutination) also typed these isolates as 33F.

Following investigation of the 33F-1 *cps* locus, it was evident that not only did all ten isolates lack *wcjE*, the locus contained *wcyO* at this position. The *wcyO* gene encodes an acetyltransferase and mediates the same modification as *wcjE* (6-O-acetylation of galactose) [21]. The *wcyO* open reading frame from all 33F-1 isolates contained a frameshift mutation. The *wcyO* gene in 33F-1 pneumococci from Fiji had a single T insertion whereas this gene in isolates from Mongolia contained a single A deletion (Fig. 1). These frameshift mutations were also confirmed by Sanger sequencing and were not present in traditional *wcyO*-containing isolates (serotypes 34 and 39) from Fiji (Supplementary Fig. S1).

**Figure 1.**
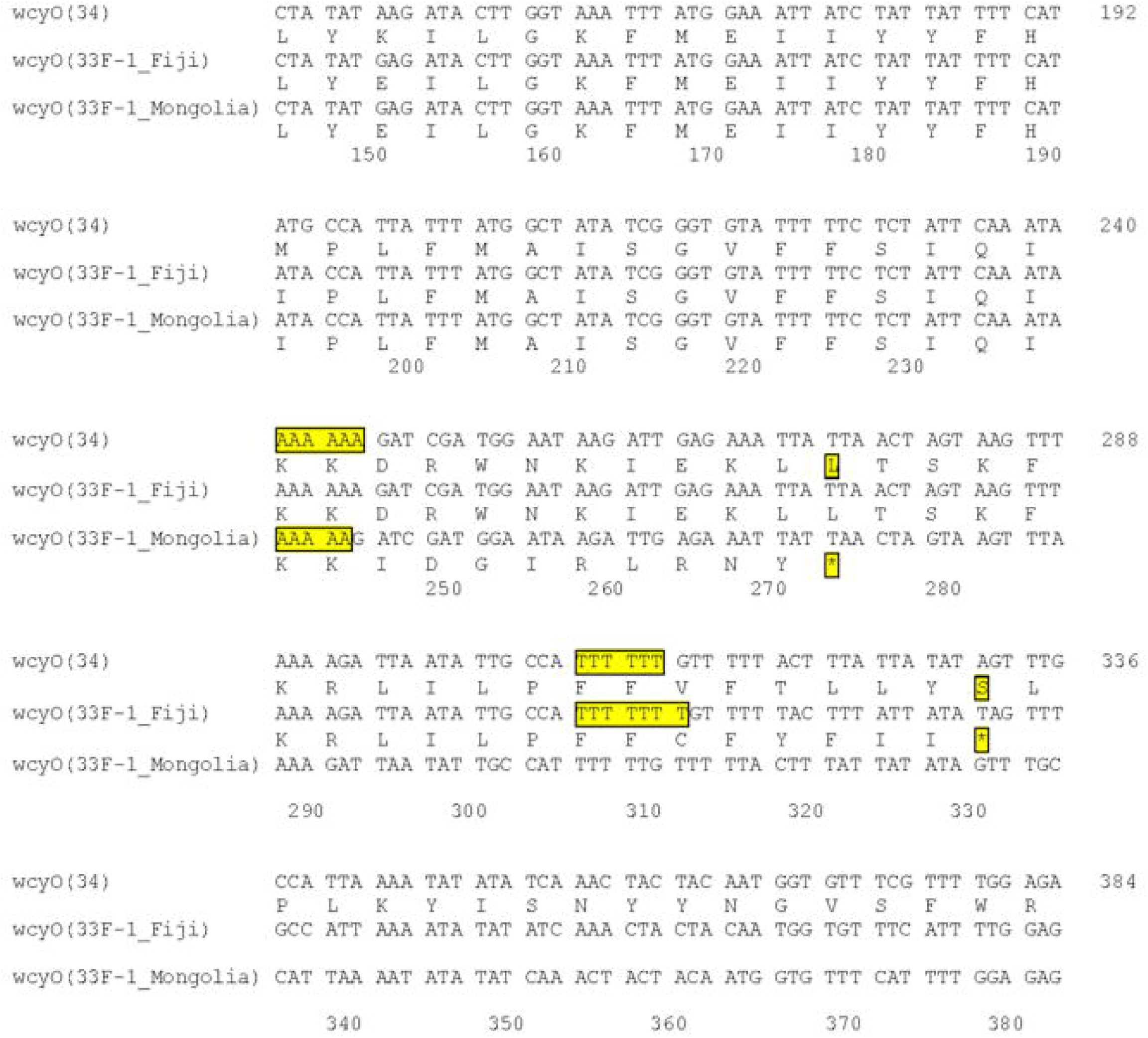
Comparison of the *wcyO* open reading frames of 33F-1 sequences to a representative serotype 34 sequence. Only a selected portion of the DNA sequence is shown. Numbers refer to the position number in the serotype 34 sequence with an in-frame *wcyO* gene.

In addition to the differences in *wcjE* and *wcyO*, microarray detected some divergence in other genes in the 33F-1 *cps* locus compared to the reference 33F sequence. To gain a better understanding of the relationships of the 33F-1 *cps* genes to homologues from other serotypes we performed phylogenetic analyses for each gene. In support of the pairwise alignments (Supplementary Table S1), the 33F-1 *wciB, wciD, wciE, wciF, wzy* and *wzx* genes clustered with 37/33A/33F sequences (Fig. 2F-K). In contrast, 33F-1 *wzg, wzh, wzd, wze* and *wchA* clustered with serotype 33B sequences (Fig. 2A-D), *wciG* with serotype 37 (Fig. 2L), *glf* with serotypes 34 and 39 (Fig. 2M) and *wcyO* with 33C, 34 and 39 (Fig. 2N). All branches had strong statistical support (>85% bootstrap score from 1000 replicates for all genes, except *wze* with a 67% bootstrap score for the 33F-1/33B branch).

**Figure 2.**
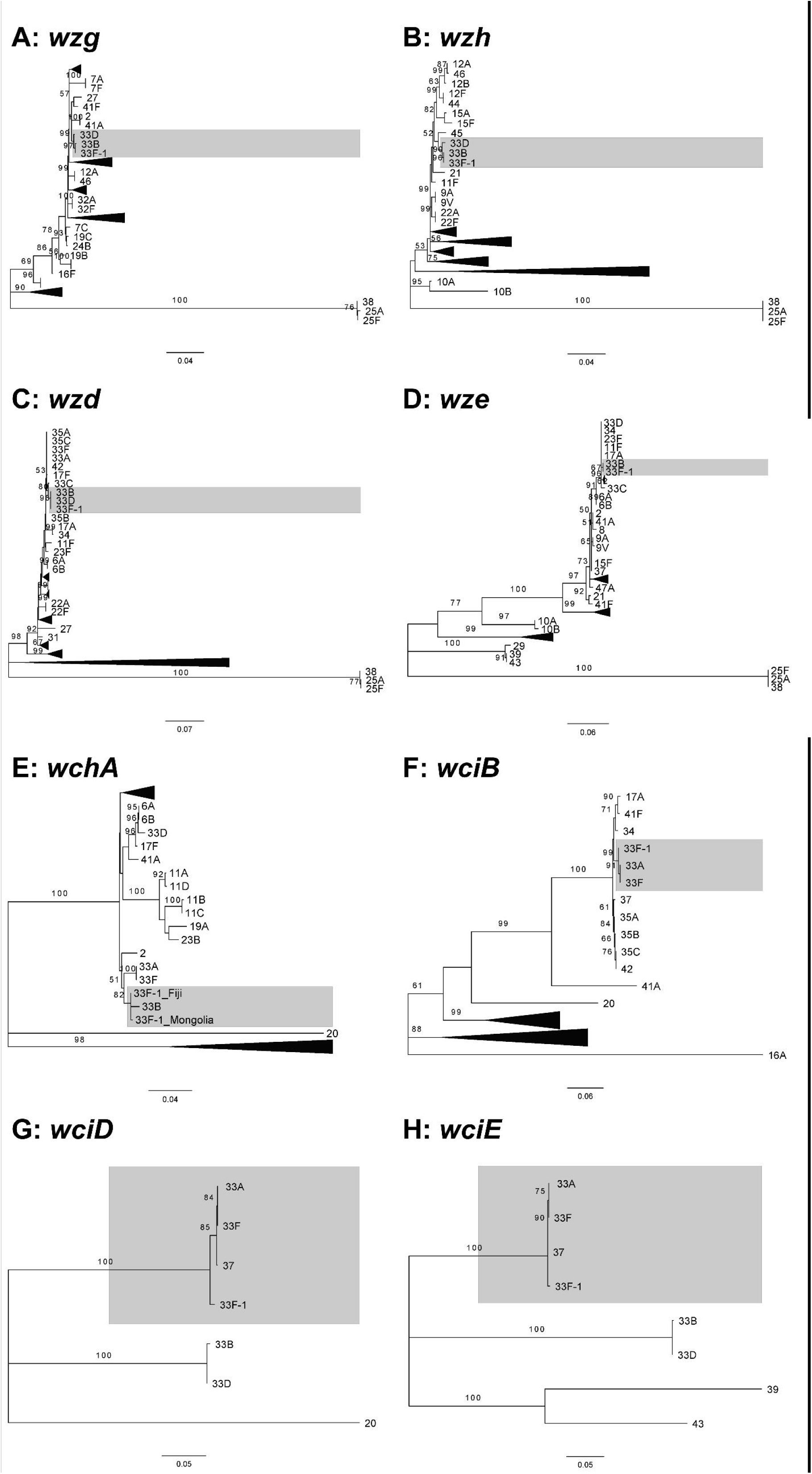
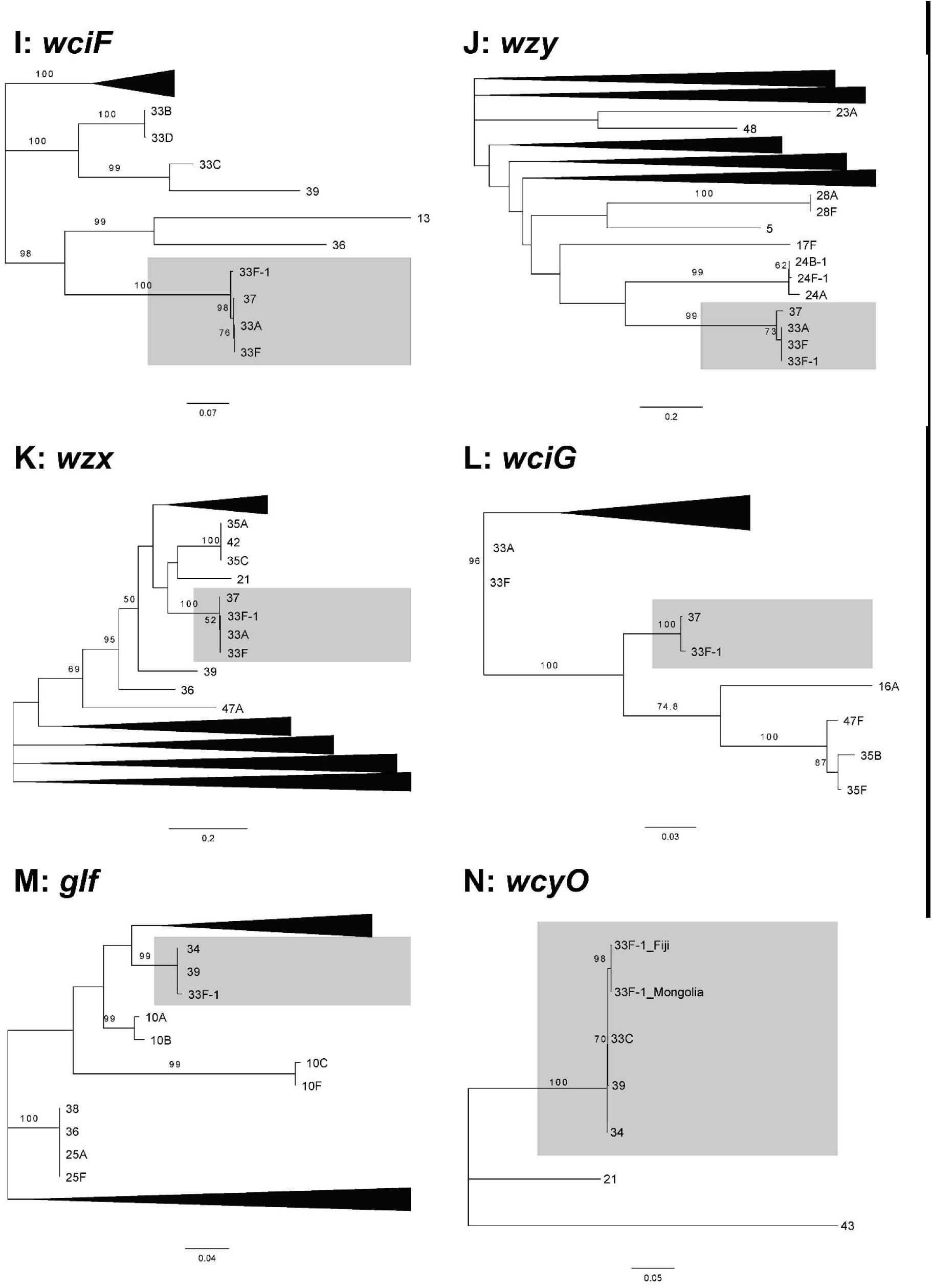
Maximum likelihood phylogenetic trees of 33F-1 *cps* genes with homologues from all other serotypes. As all genes except *wchA* and *wcyO* were identical in all 33F-1 isolates only one sequence is included as a representative. Un-collapsed trees are provided in Supplementary Figure S2. Tree for *wciC* is not shown as this gene is only present in serotypes 33F, 33A and 37, which all have over 98% DNA sequence identity to the 33F-1 sequence. DNA sequences were aligned using MUSCLE and trees were constructed using the Tamura-Nei model in MEGA 7. Only bootstrap values above 50% are shown.

## Discussion

Pneumococcus is a highly successful pathogen, in part due to the high level of capsule diversity, resulting in over 90 serotypes each with unique antigenic properties. Even small differences in the *cps* locus can have biologically relevant consequences. Serotypes 33F and 33A have the same *cps* locus, except that 33F has a *wcjE* gene containing a frameshift mutation rendering it non-functional [22]. Using DNA microarray, we identified a high degree of genetic divergence in the capsule DNA sequence of some serotypes in Fiji and Mongolia. We characterized a serotype 33F variant (33F-1) that has the same genes as the canonical 33F and 33A *cps* loci, except it possesses *wcyO* instead of *wcjE*. Interestingly, in the 33F-1 variants *wcyO* is predicted to encode a truncated protein due to a frameshift mutation. These frameshift mutations suggest a loss of 6-O-acetylation in 33F-1 capsular polysaccharide as the truncated protein would unlikely be functional. Interestingly, the same variant has been simultaneously identified in the Global Pneumococcal Sequencing Project in other countries (van Tonder et al, unpublished), demonstrating 33F-1 pneumococci are not restricted to Fiji and Mongolia.

The frameshift mutations in isolates from Fiji and Mongolia have both occurred within homopolymeric regions (Fig. 1, Supplementary Fig. S1). Such regions are prone to slipped-strand mispairing, whereby errors made during DNA replication can result in the insertion or deletion of a nucleotide [23]. We postulate that the frameshift mutations in the 33F-1 *wcyO* genes are the result of slipped-strand mispairing events.

This is the first report identifying the *wcyO* acetyltransferase gene in the 33F *cps* locus, and it is also the first report of a naturally occurring frameshifted allele of *wcyO*. The fact that the mutation type and location differ between isolates from Fiji and Mongolia demonstrates this mutation event has occurred on at least two independent occasions. Whether the mutation of *wcyO* is due to selective pressure to inactivate a disadvantageous gene or due to a lack of selective advantage to maintain it remains to be investigated. Previously, mutations have been identified in other pneumococcal capsule acetyltransferase genes including *wciG* [24] and *wcjE* [22,25,26]. Serotype 11E, which lacks WcjE-mediated acetylation can evade opsonophagocytosis more efficiently compared to 11A (which possess WcjE-mediated acetylation) [25]. Pneumococci expressing 33F capsules, which lack WcjE-mediated acetylation, exhibit enhanced survival during drying compared to serotype 33A (with intact WcjE-mediated acetylation) [27]. Laboratory constructed *wciG* mutants in serogroup 33 isolates were more susceptible to opsonophagocytosis, and displayed increased adherence and biofilm formation [27]. It is plausible that mutation of *wcyO* in the 33F-1 pneumococci may serve a similar purpose, however this requires further investigation.

Within the 33F-1 *cps* locus we identified 7/15 genes that exhibit higher DNA sequence similarity to homologues from other serotypes rather than 33F. Both *glf* and *wcyO* are similar to sequences from serotypes 34 and 39 (and 33C for *wcyO)* (Fig. 2M and 2N) and *wzg, wzh, wzd, wze* and *wchA* similar to sequences from 33B (Fig. 2A-E). Recombination of the pneumococcal capsule genes resulting in mosaic *cps* loci such as that of 33F-1 have been reported previously [28,29]. Although it is difficult to infer the direction of horizontal transfer of these genes, the mosaic nature of the 33F-1 *cps* locus would suggest an ancestral 33A/F *cps* locus was the recipient of these genes.

Interestingly, a serogroup 33 related *cps* locus has been identified in *Streptococcus oralis* subsp. *tigurinus* strain Az_3a [30]. This *cps* locus possessed the same genes as the 33F-1 locus with variable DNA identity (<77% with the 33F-1 *wzg, wzh, wzd, wze, wchA* and *wciB* genes, >96% for *wciC, wciD, wciE, wciF* and *wzy* genes, and 85–90% for *wzx, wciG* and *glf* genes, Supplementary Table S1). The higher DNA identity of 33F-1 *cps* genes with homologues from other pneumococcal serotypes suggests the Az_3a *cps* locus may have evolved independently of the 33F-1 locus. In contrast to 33F-1, the *wcyO* gene in Az_3a is in frame and most similar to the pneumococcal serotype 21 homologue (DNA identity 86.8% with serotype 21 *wcyO* compared to 74.5% with 33F-1 *wcyO)*. The existence of a divergent 33F-1 *cps* locus with a functional *wcyO* raises interesting questions around why this gene has been inactivated in 33F-1 pneumococci but remains intact in a non-pneumococcal streptococcal species.

This study describes a novel genetic basis for pneumococcal serotype 33F. Serotype 33F is a replacing serotype in invasive disease following vaccine introduction [5–7]. The public health importance of 33F is reflected in that it has been included in two upcoming vaccine formulations (PCV15 and PCV24) [8]. In addition, there is increasing popularity in molecular serotyping approaches and it is therefore important to identify genetic variants, which have the potential to impact serotyping results. This is particularly important for the implementation of such methods in LMICs, where there is limited understanding of the pneumococcal *cps* loci. The data gained from this study will be used to update genetic typing tools for more accurate typing of serotype 33F in LMICs.

## Acknowledgments

We thank the participants, their families and villages; Fiji Ministry of Health and Medical Services and the Ministry of Health in Mongolia. We also thank all study staff involved in recruitment, swab collection and laboratory analyses, including Suuri Bujinlham, Tupou Ratu, Silivia Mantanitobua, Evelyn Tuivaga and Mere Guanivalu. This study was supported by the Bill & Melinda Gates Foundation (OPP1126272, OPP1084341 and OPP115490); Gavi, the Vaccine Alliance; and the Department of Foreign Affairs and Trade of the Australian Government and Fiji Health Sector Support Program (FHSSP). Catherine Satzke holds a NHMRC Career Development Fellowship and a veski Inspiring Women Fellowship. Sam Manna received a Robert Austrian Research Award in Pneumococcal Vaccinology funded by Pfizer. This work was also supported by the Victorian Government's Operational Infrastructure Support Program.

## Transparency declaration

The authors declare that they have no conflicts of interest relevant to this article.

